## Supplementary figures and images for "A population-level invasion by transposable elements triggers genome expansion in a fungal pathogen"

### F1-Suppl1

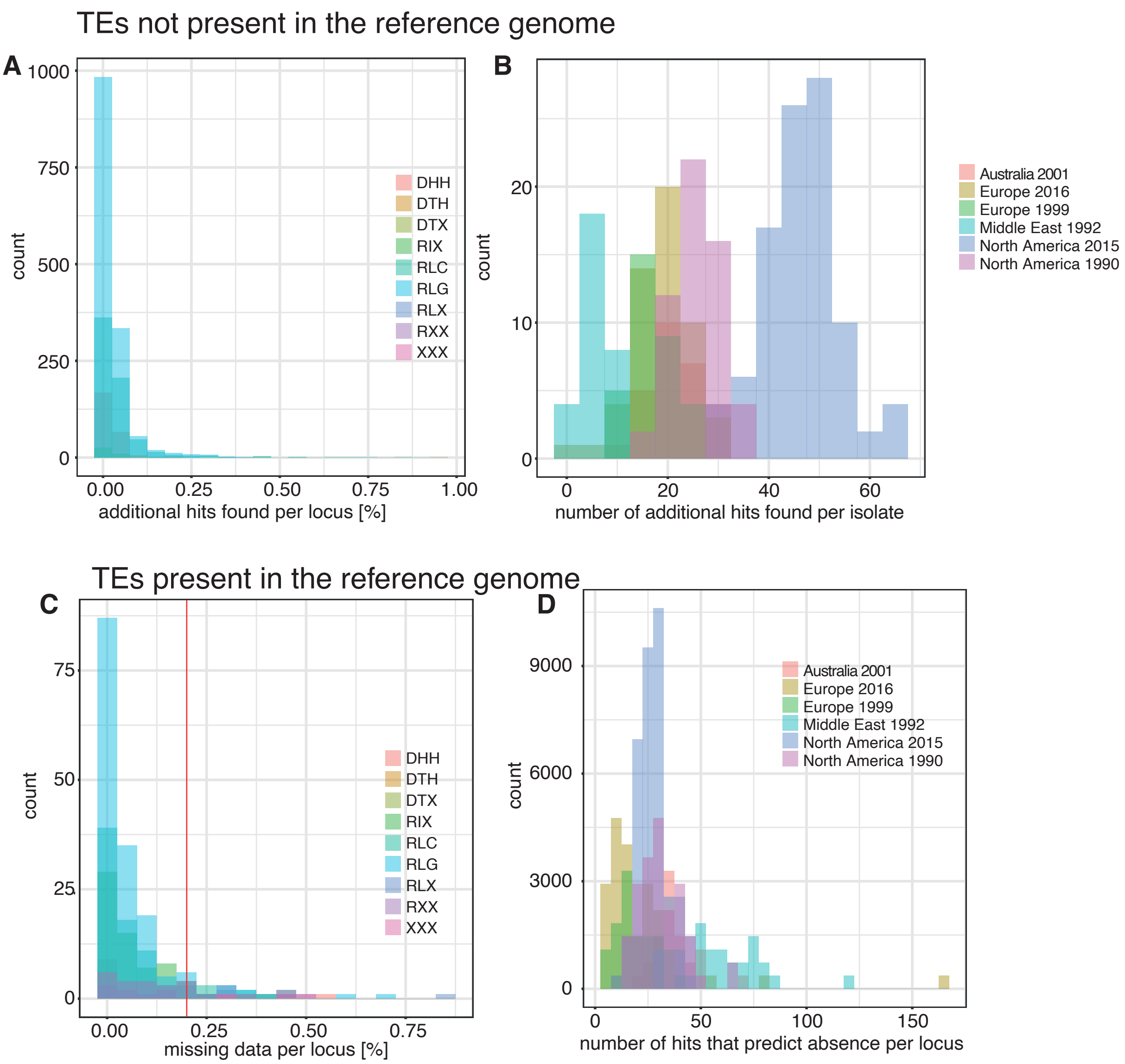

### F1-Suppl3

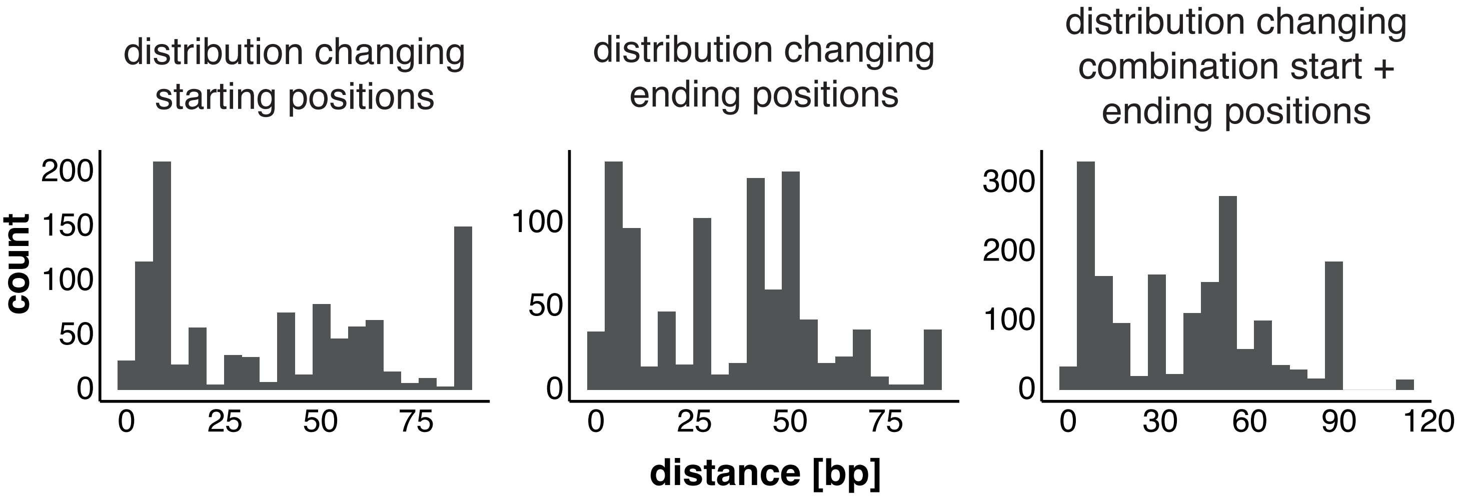

### F1-Suppl4

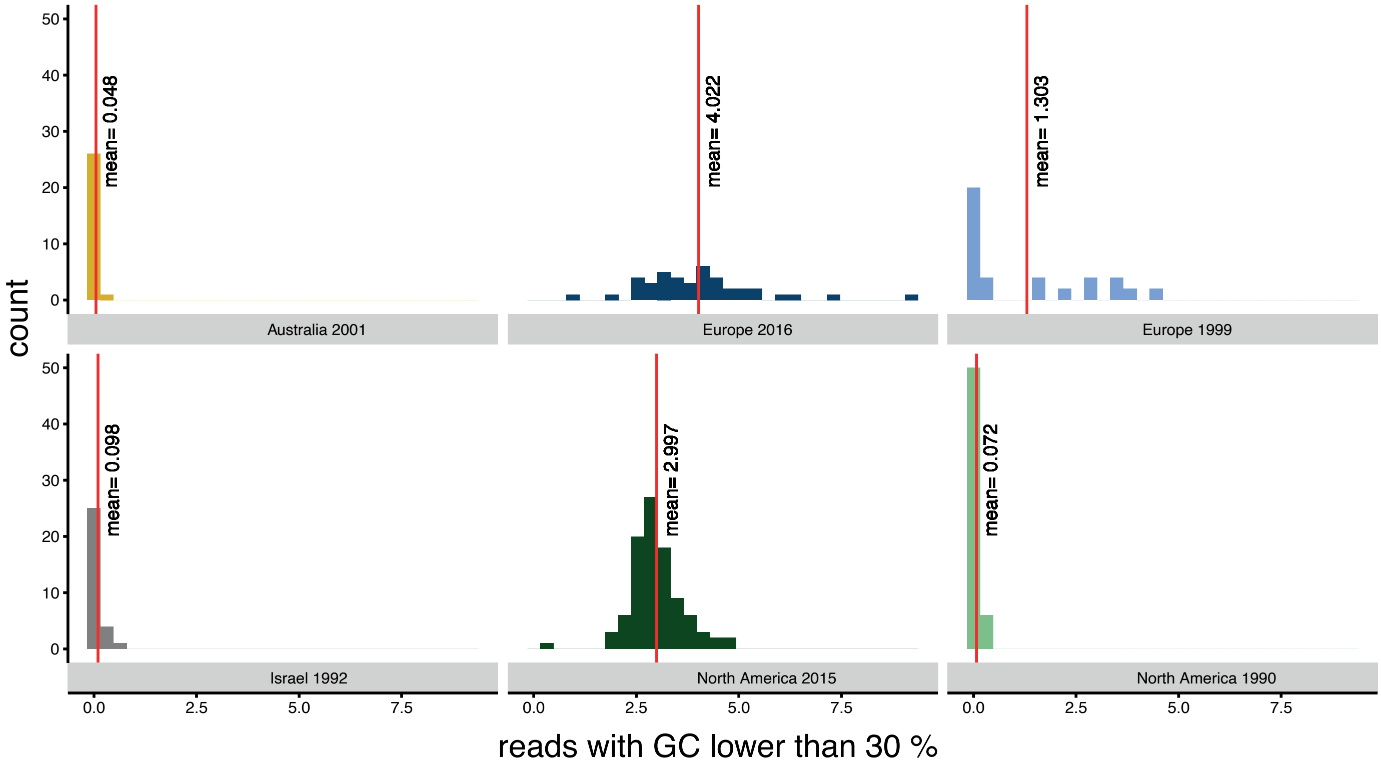

### F2-Suppl2

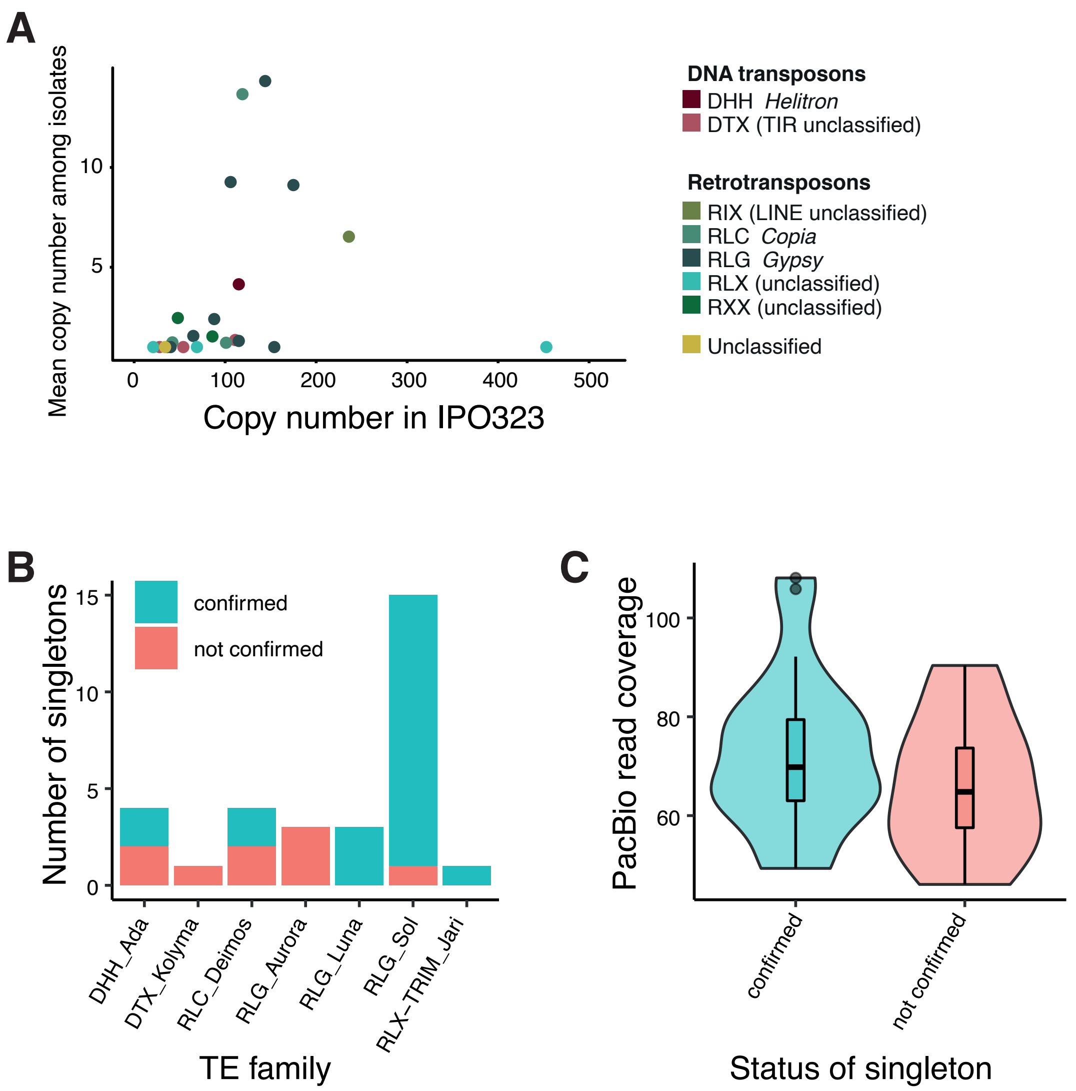

### F2-Suppl5

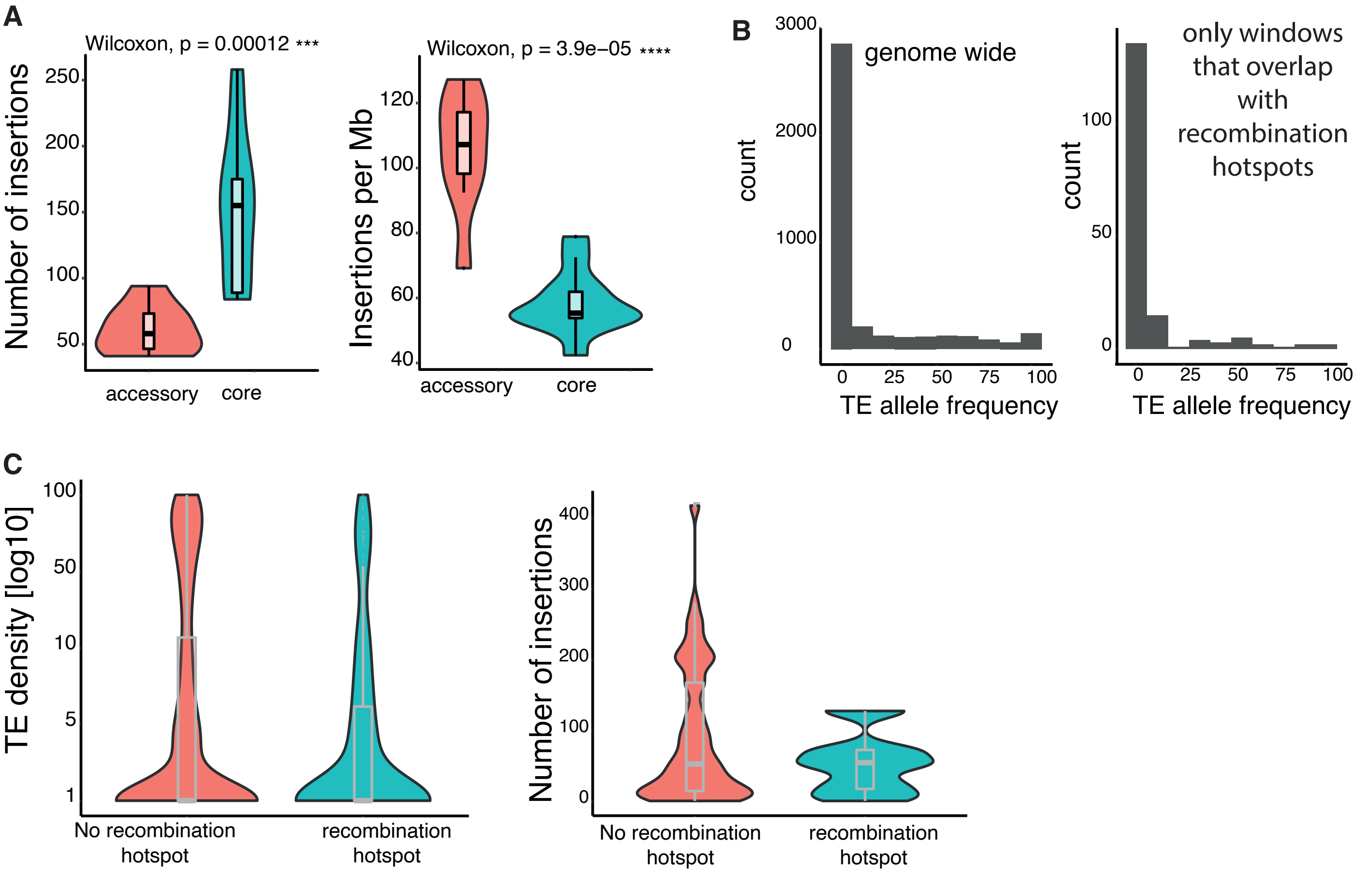

### F2-Suppl6

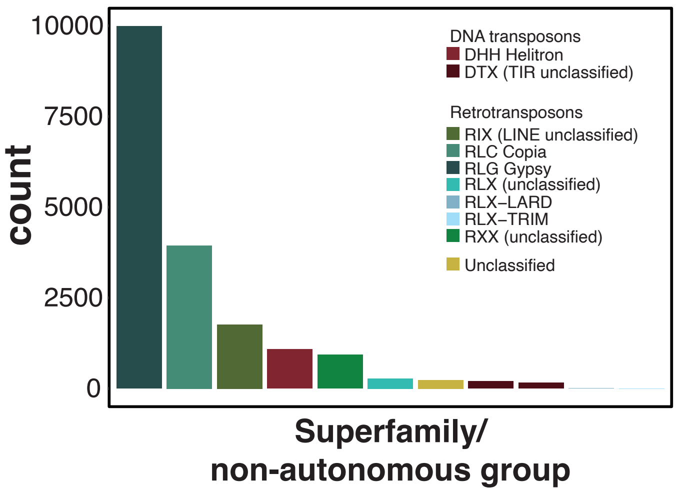

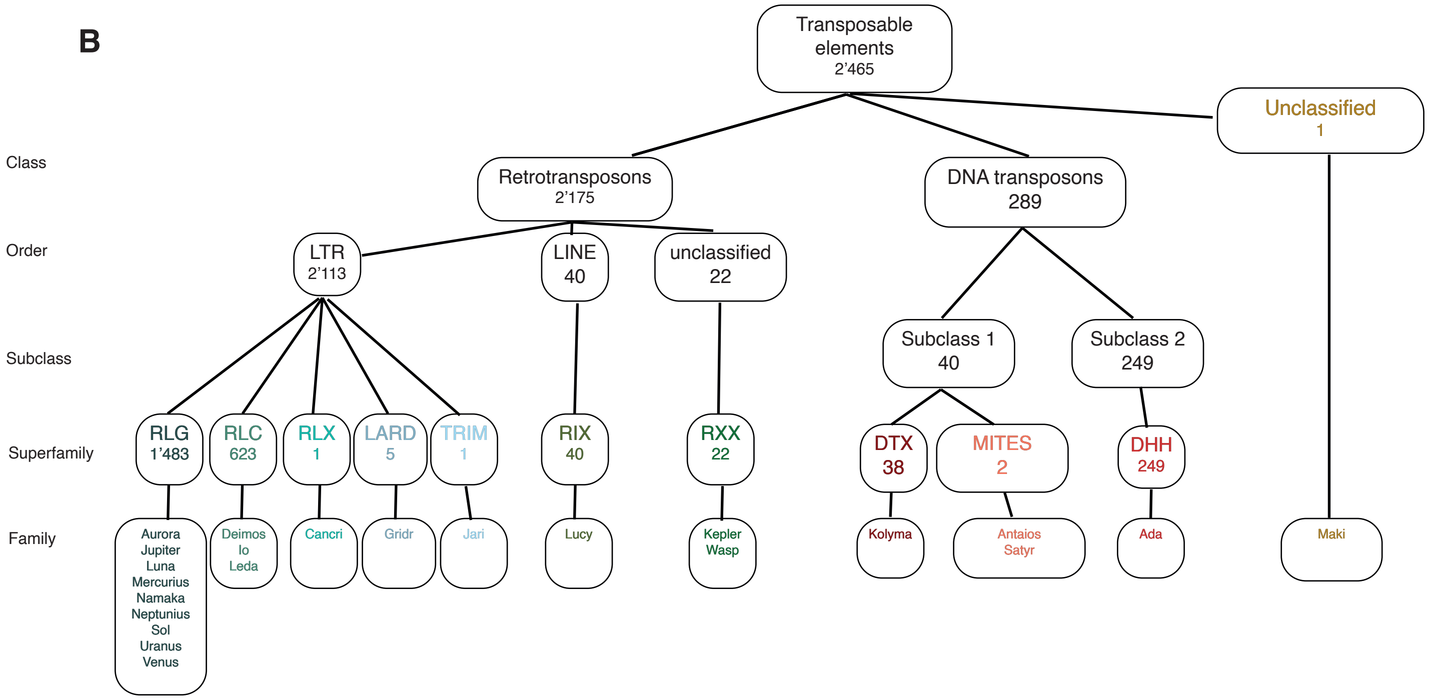

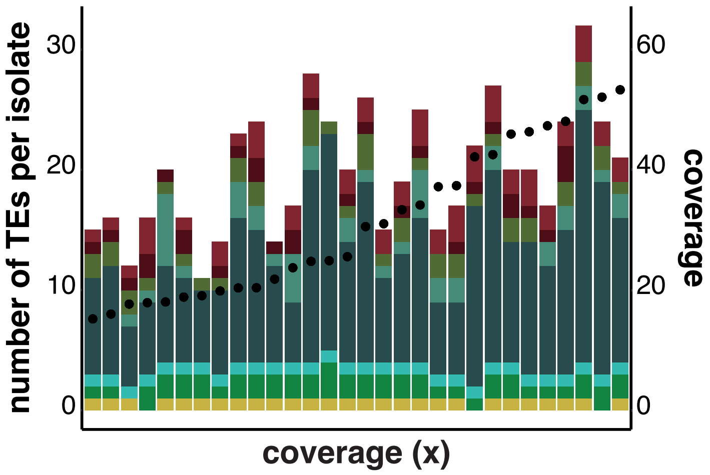

### F3-Suppl1

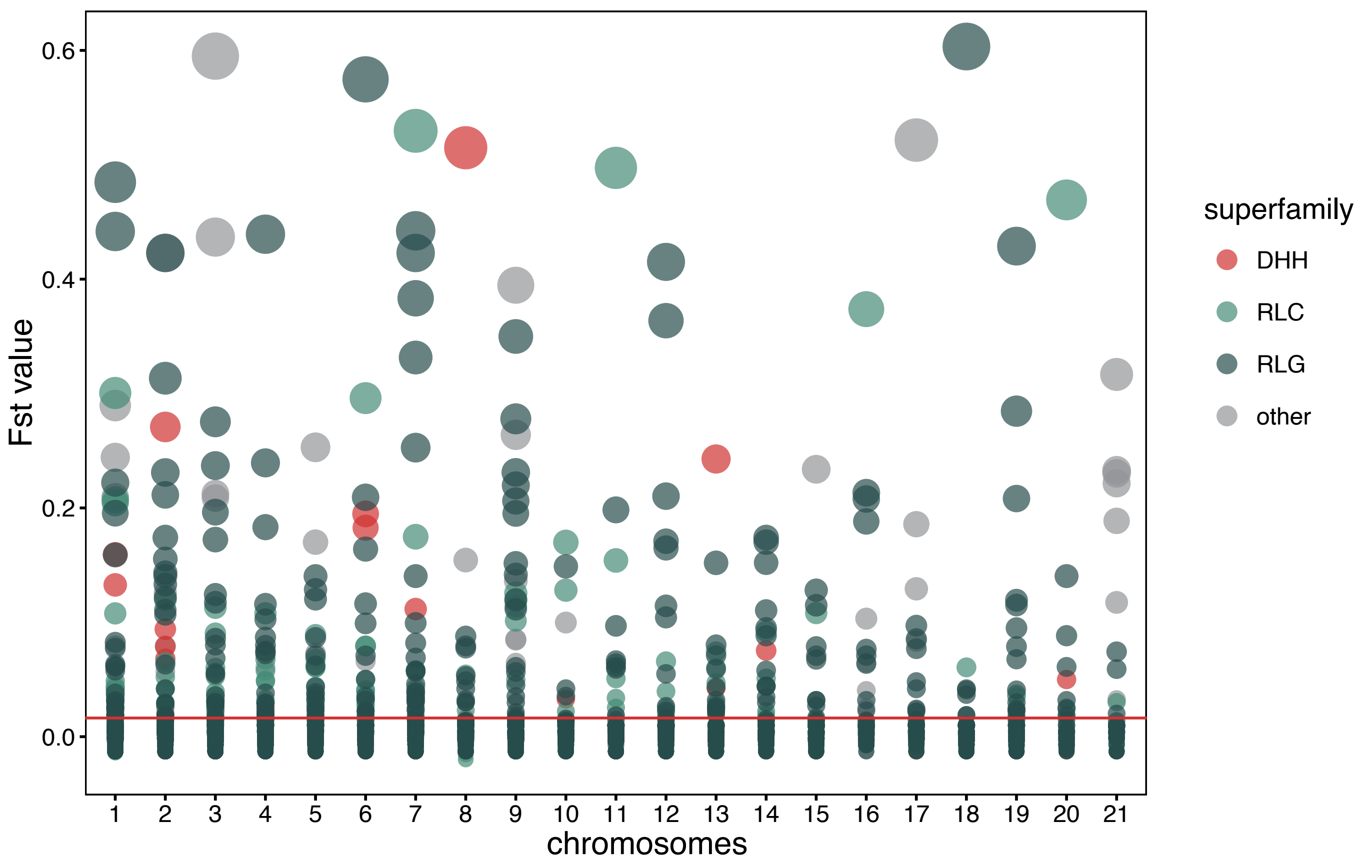

### F4-Suppl2

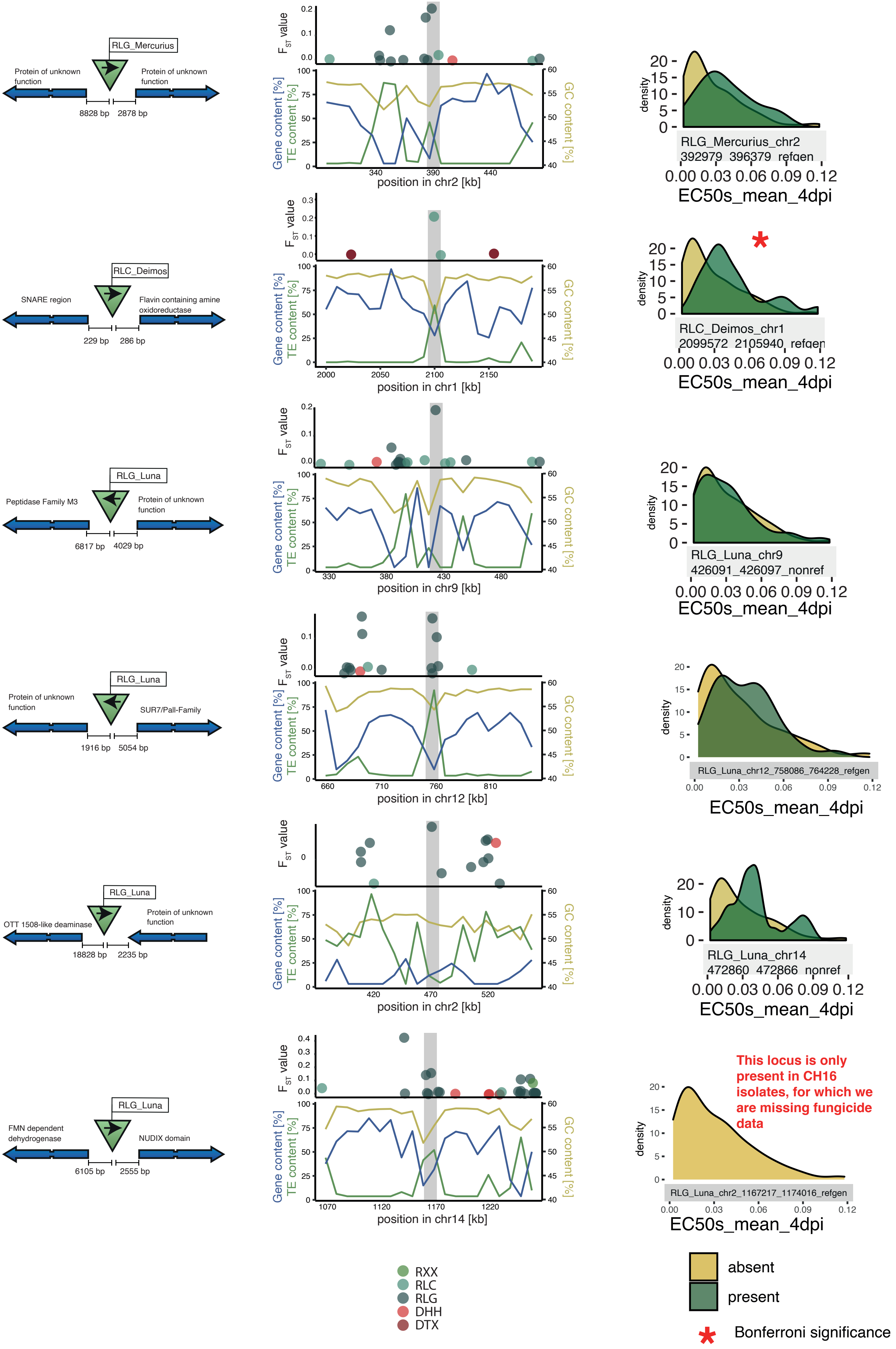

### F6-Suppl1

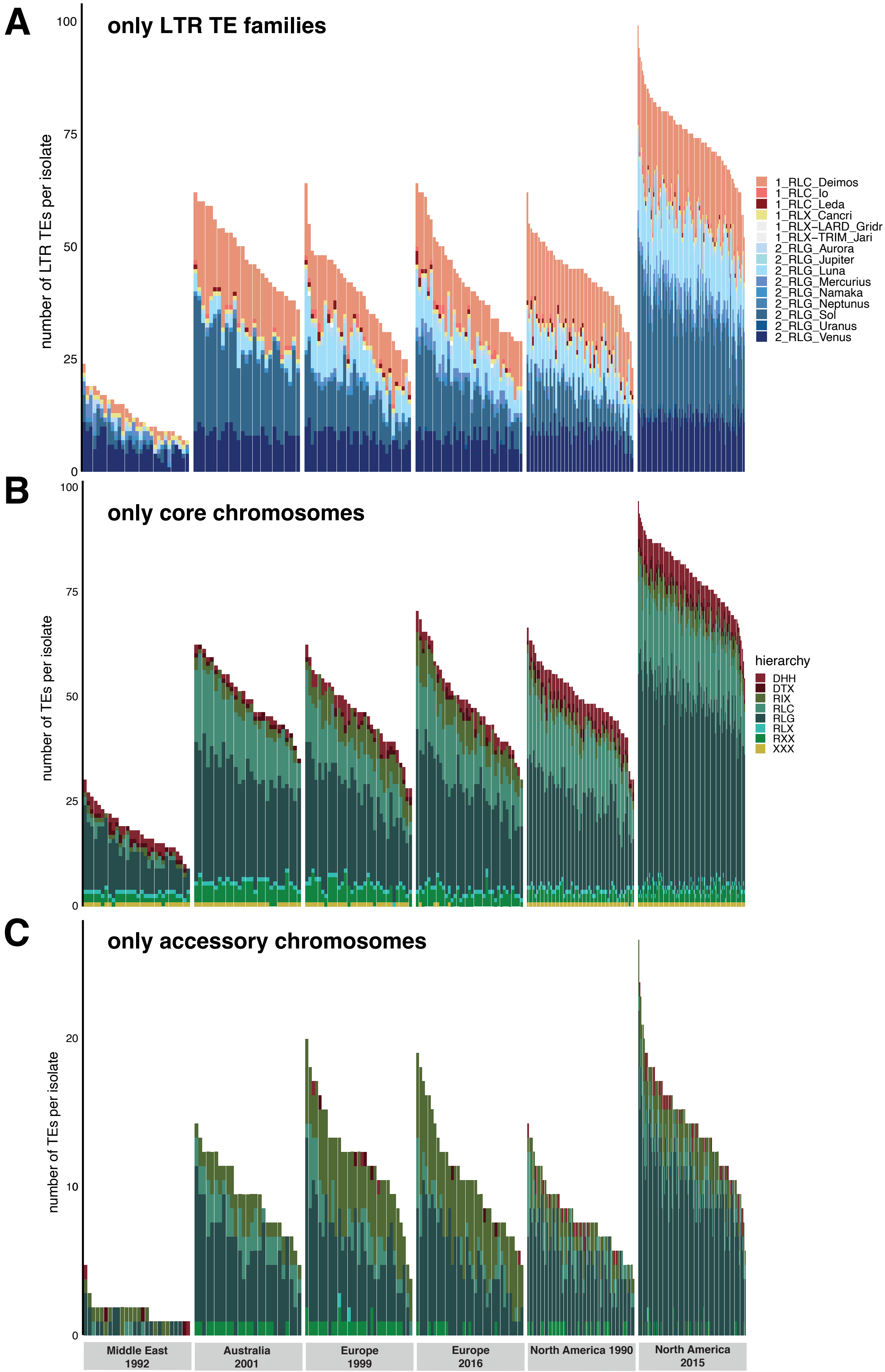

### F6-Suppl3

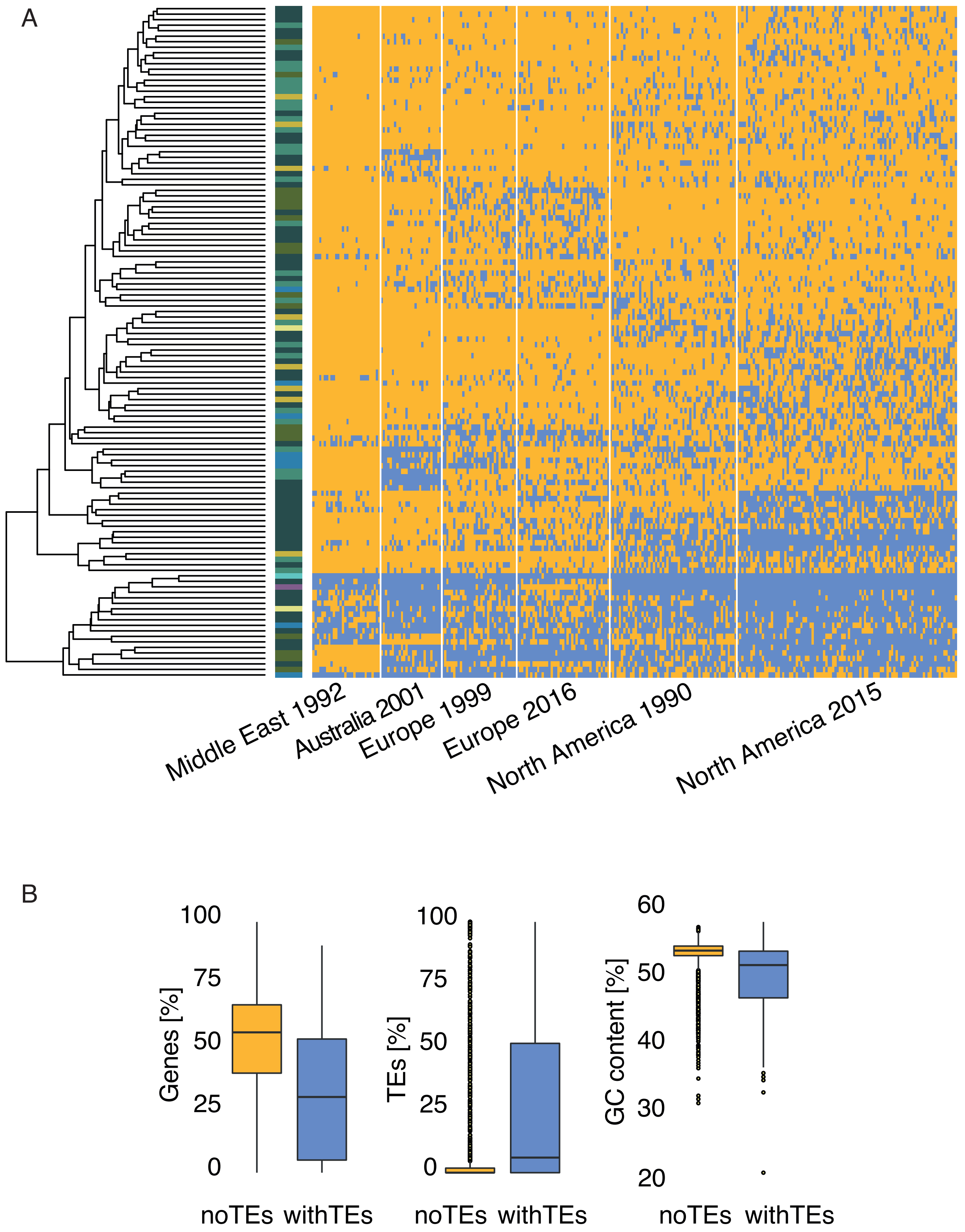

### F7-Suppl1

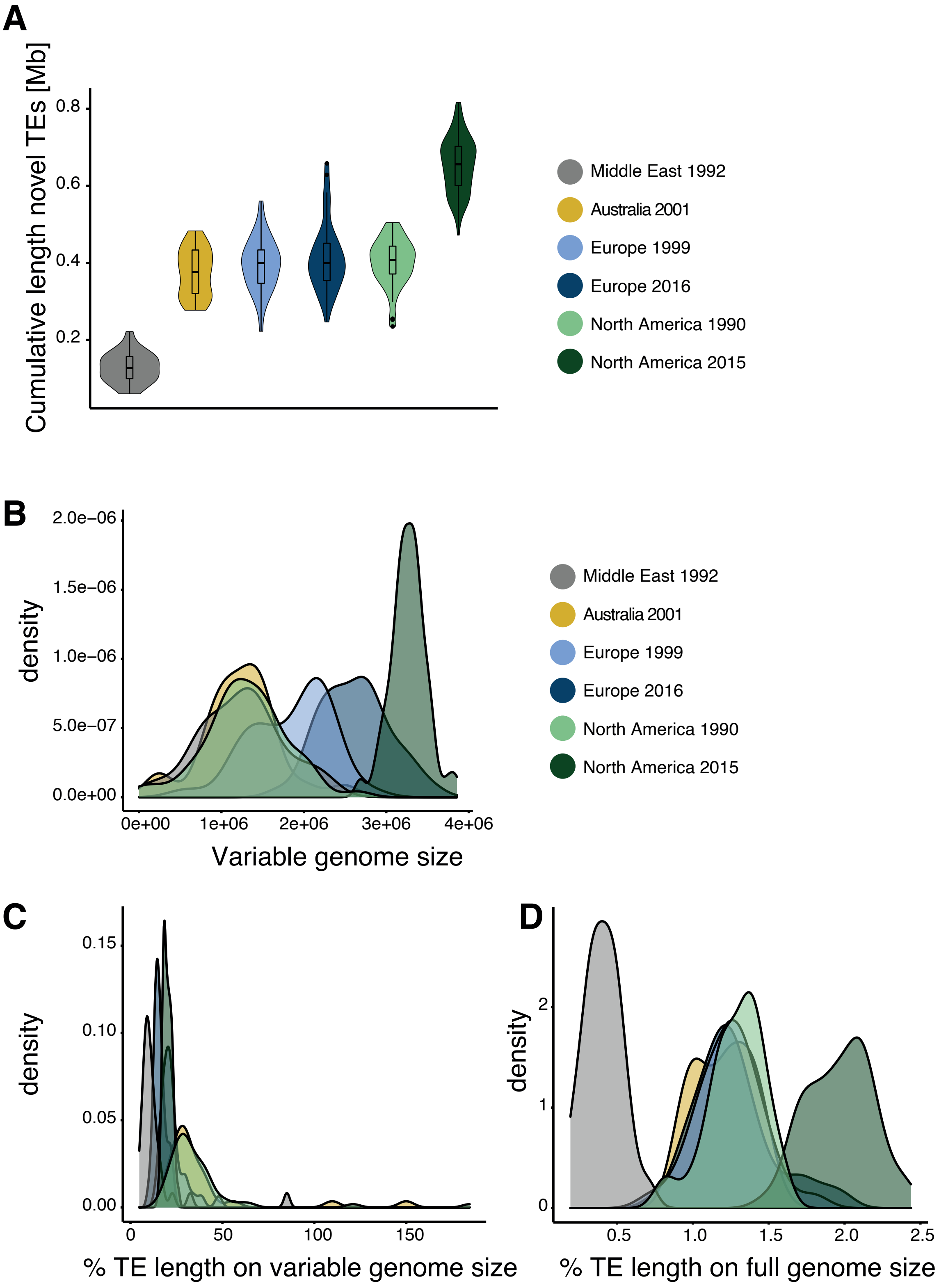
